# Fully Automated MRI-based Analysis of the Locus Coeruleus in Aging and Alzheimer’s Disease Dementia using ELSI-Net

**DOI:** 10.1101/2024.07.26.605356

**Authors:** Max Dünnwald, Friedrich Krohn, Alessandro Sciarra, Mousumi Sarkar, Anja Schneider, Klaus Fliessbach, Okka Kimmich, Frank Jessen, Ayda Rostamzadeh, Wenzel Glanz, Enise I. Incesoy, Stefan Teipel, Ingo Kilimann, Doreen Goerss, Annika Spottke, Johanna Brustkern, Michael T. Heneka, Frederic Brosseron, Falk Lüsebrink, Dorothea Hämmerer, Emrah Düzel, Klaus Tönnies, Steffen Oeltze-Jafra, Matthew J. Betts

## Abstract

**INTRODUCTION:** The Locus Coeruleus (LC) is linked to the development and pathophysiology of neurodegenerative diseases such as Alzheimer’s Disease (AD). Magnetic Resonance Imaging based LC features have shown potential to assess LC integrity in vivo.

**METHODS:** We present a Deep Learning based LC segmentation and feature extraction method: ELSI-Net and apply it to healthy aging and AD dementia datasets. Agreement to expert raters and previously published LC atlases were assessed. We aimed to reproduce previously reported differences in LC integrity in aging and AD dementia and correlate extracted features to cerebrospinal fluid (CSF) biomarkers of AD pathology.

**RESULTS:** ELSI-Net demonstrated high agreement to expert raters and published atlases. Previously reported group differences in LC integrity were detected and correlations to CSF biomarkers were found.

**DISCUSSION:** Although we found excellent performance, further evaluations on more diverse datasets from clinical cohorts are required for a conclusive assessment of ELSI-Nets general applicability.

**Highlights:**

- thorough evaluation of a fully automatic LC segmentation method termed ELSI-Net in aging and AD dementia
- ELSI-Net outperforms previous work and shows high agreement with manual ratings and previously published LC atlases
- ELSI-Net replicates previously shown LC group differences in aging and AD
- ELSI-Net’s LC volume correlates with CSF biomarkers of AD pathology

**RESEARCH IN CONTEXT:**

1. **Systematic Review:** The authors reviewed the literature using traditional sources (e.g. Pubmed, Google Scholar). Although there are several publications introducing semi-automatic methods for LC segmentation, the application of Deep Learning methods is underexplored. To the best of our knowledge, this is the first paper using a Deep Learning based approach for automated LC segmentation in AD dementia.
2. **Interpretation:** Our work introduces and evaluates an improved automatic, Deep Learning based LC segmentation and analysis approach. The results suggest a very high potential for practical applicability, e.g. in large-scale clinical studies for neurodegenerative diseases.
3. **Future Directions:** ELSI-Net can be used to assess LC integrity on large- or small-scale studies in Alzheimer’s Disease dementia. To ensure robust performance, ELSI-Net should be further evaluated in larger, more diverse datasets comprising varying LC MRI protocols and clinical populations.

## 1 Background

The Locus Coeruleus (LC) is a small brainstem nucleus with about 50.000 pigmented neurons,[1] but projects to almost all major brain regions and is the primary source of noradrenaline in the brain. It has been identified as one of the earliest brain structures to be affected in Alzheimer’s Disease (AD)[2] and has been linked to cognitive decline in healthy aging and progression of AD.[3] Owing to the early tau aggregation in the LC, assessing the integrity of the LC using structural MRI may be a suitable tool for obtaining pathophysiological insights in vivo.[4, 5] It presents an opportunity to gain insights into cognitive and behavioral symptoms instrumental for developing effective treatments[1], improve our understanding of AD pathogenesis and facilitate the development of disease-modifying noradrenergic drugs.[6]

So-called neuromelanin-sensitve Magnetic Resonance Imaging (MRI) techniques permit the in vivo visualization of the LC by exploiting the magnetic properties of its neuromelanin pigmented neurons although the exact mechanisms remain unclear.[4, 7] The hyperintense regions appearing in the MRI acquisitions were shown to correspond to LC properties observed in post-mortem studies, i.e. with respect to anatomical position and dimensions, LC cell density[8] and to correspond with age-related increases in neuromelanin.[9] Furthermore, associations between LC MRI contrast and AD biomarkers[10] as well as cognitive decline in health and disease have also been observed[5] suggesting neuromelanin-sensitive MRI may be suitable for assessing LC integrity.

A reliable extraction of in vivo LC MRI biomarkers requires a robust segmentation approach. The small size and cylindrical shape of the LC (approximately 2mm in diameter[11]) together with the comparatively coarse resolution of MRI acquisitions limits the reliability of LC integrity measures. Although reasonable compromises can be found,[4] LC MRI acquisitions are characterized by low signal-to-noise ratios and ambiguous structural boundaries posing challenges for segmentation approaches. This is evident by particularly low inter-rater agreements between manual raters of 0.499,[12] 0.54 to 0.64[13] and 0.67 Dice Similarity Coefficient (DSC)[14] as reported in the literature.

Initially, a broad majority of studies investigating the LC using neuromelanin-sensitive MRI relied on manual segmentations carried out by expert raters.[15] They require considerable amounts of manual labour and time of trained experts. In recent years, (semi-)automatic approaches, that reduce the need for manual intervention, have become more common than purely manual segmentation. Many studies have used atlas or template-based segmentation approaches.[8, 12, 16, 17] Another type of algorithms employs the template registration as a first step to obtain a search space on which further operations, usually related to peak intensity extraction of certain rostrocaudal subparts or slices of the LC, are applied.[18, 19] Although only a subset of the LC voxels are obtained, promising results using intensity-based features have been shown with respect to reproducing known cohort effects, such as structural degeneration,[19] associations to cognition or both.[20] These methods require manual corrections in some cases as the performance relies on successful registration with a high precision. Through the remaining manual steps, rater bias may still influence the segmentation of these methods. Automatic LC segmentation algorithms such as the approach presented here, may facilitate using LC imaging in large-scale studies by removing the need for multiple experts to invest time and manual labour on LC segmentation and introduce more objectivity by removing human bias. Our group was the first to develop a Deep Learning-based LC segmentation approach.[21] Its convolutional neural networks process the MRI acquisitions inherently faster than registration-based and manual methods and have shown higher objectivity by incorporating multiple experts’ knowledge into the training.[14] Our pipeline comprises all steps from the LC segmentation to the reference region generation and the feature extraction. We do not manually correct the segmentations prior to our analyses and evaluations.

In the work presented here, we show an improved fully automatic LC segmentation pipeline that further increases the performance compared to our previous approach and assess its practical usability in various ways on subjects of aging and AD dementia.

## 2 Material and Methods

We apply an improved version of a recently proposed fully automatic LC analysis method[14] to two different datasets comprising MRI acquisitions from healthy aging and Alzheimer’s disease dementia. The results are compared to manual expert segmentations and their features.

### 2.1 Datasets

The two datasets share the same acquisition protocol: They comprise T_1_-weighted Fast Low-Angle Shot (FLASH) 3 Tesla MRI scans (5.56ms echo time, 20ms repetition time, 23° flip angle, 130Hz/pixel bandwidth, 7/8 partial Fourier, 13:50min scan time) with an isotropic resolution of 0.75mm. The image data was upsampled using a sinc filter to achieve an isotropic voxel size of 0.375mm and then bias-field-corrected as previously described.[9] Our Healthy Aging Dataset (HAD)[9] comprised 82 healthy subjects. There were 25 younger (22-30 years old; 13 male) and 57 older subjects (61-80 years old, 19 male). We also analyzed neuromelanin-sensitive MRI data from the DZNE Longitudinal Cognitive Impairment and Dementia study (DELCODE).[22] This comprised 188 subjects: 68 healthy elderly adults, 22 relatives of individuals with AD, 61 subjects with subjective cognitive decline (SCD), 26 with mild cognitive impairment (MCI) and 11 with AD dementia. For our experiments, we combine the healthy elderly adults and relatives of AD subjects to one group of healthy controls. The age of the subjects ranges from 60 to 87 years (ca. 69 on average, 102 females). The neuromelanin-sensitive MRI data has been acquired at four different sites across Germany: Magdeburg, Rostock, Bonn and Berlin. We refer the reader to Betts et al.[9] (for HAD) and Jessen et al.[22] for further details on the full DELCODE cohort.

### 2.2 Segmentation Methods

#### 2.2.1 Manual Expert Segmentation

Both of our trained expert raters (M. Betts, referred to as Rater 1 (R1) and M. Sarkar, referred to as Rater 2 (R2)) manually segmented all LCs in the HAD. For the DELCODE dataset, R1 manually segmented 108 subjects and R2 segmented the remaining 80 subjects.

The raters delineated the LC using ITK-SNAP[23] as previously described.[9] Briefly, the segmentation was performed on the axial slices starting at the most dorsal to ventral portion of the LC while limiting the rostrocaudal space for segmentation to slices between the inferior boundary of the interpeduncular fossa at the level of the inferior colliculus and the superior cerebellar peduncle. (for reference see[9]).

#### 2.2.2 Ensemble-based Locus Coeruleus Segmentation Network (ELSI-Net)

Our previously published approach[14] for fully automatic LC segmentation was further improved and used in this work. Figure 1 shows a schematic overview of the method comprising two fundamental steps that are realized using two almost identical 3D U-Net based[24] convolutional neural networks: initial LC localization followed by segmentation on an extracted patch containing only the LC and its immediate vicinity. To train and evaluate our model, we ran a 3×5-fold nested crossvalidation which splits the subjects of the HAD dataset just as in our previous work[14] making the results directly comparable. A final training was conducted splitting the HAD in 5 equally sized subsets to obtain the 5 nets (each trained with one of these subsets as validation set and the combined rest as training set) that were used for the application to DELCODE. There was no fine-tuning or retraining with DELCODE subjects and they have not been used for either, the training or validation sets (for early stopping), so that it can be seen entirely as a test set.

**Figure 1:**
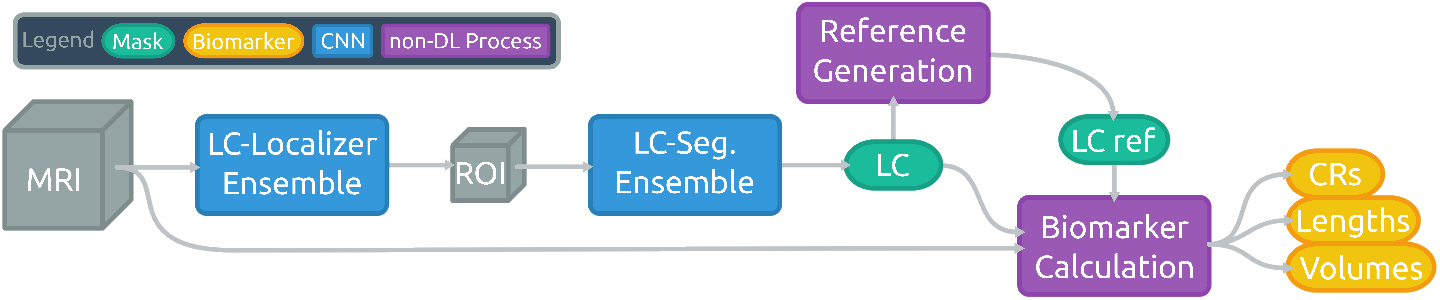
Schematic illustration of the ELSI-Net pipeline for automated LC analysis.

Three changes were introduced to the method compared to our previous work.[14] Firstly, for the application of the models, we combined the five resulting networks in an ensemble and conducted an averaging and majority vote on the different outputs to determine the final predictions for the localization and segmentation nets, respectively. This way, the final result can profit from the information obtainable from the entire training set despite the necessity for a validation set for each individual network. Second, we normalized the intensities of the extracted image patch once again prior to passing it to the segmentation network aiming to reduce the variance of the intensity range.

Third, we replaced the reference region generation relying on a sufficiently accurate Pons segmentation. Instead, we determine an LC-oriented orthonormal vector base forming a coordinate system and we calculate the average offset of the semi-automatically generated reference regions on the training set in relation to the respective LCs. They are located in the pontine tegmentum - one per hemisphere. In the application case, we determine the same coordinate system and place the reference region according to the learned offset. This approach does not require a reliable Pons segmentation or time-consuming registration procedures and is potentially more robust to head rotations incurred during acquisitions. The vectors for this LC-oriented coordinate system are determined as follows.

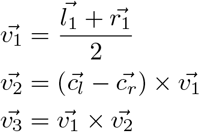

with 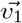 the rostrocaudal LC direction derived from the two principal components obtained from two principal component analyses on the mask voxel coordinates of the LC masks of the left 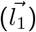 and right hemisphere 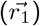, the second base vector 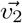, the center of mass of the left 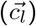 and right LC 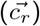 and the third base vector 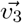.

### 2.3 Feature Extraction

The rostral LC may be particularly vulnerable in AD.[10] Hence, we extracted not only the entire LC MRI contrast ratios (CRs), but also subregional CRs, LC volume and length. Throughout this work, all reported features are bilateral, i.e. they are the average of both LC hemispheres’ features.

#### contrast ratios (CRs)

The most frequently used LC feature in the literature[15, 25] are MRI intensity ratios, that calculate the ratio of the maximum or median intensity value of the voxels in the LC mask (LC_max_ or LC_median_) to the median value of a reference region (REF_median_), positioned in the pontine tegmentum. For example, the maximum CR is defined as follows:

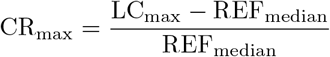

#### subregional CRs

We furthermore calculate the CRs of subregions of the LC by splitting it along its axial dimension into two and three sections equal of length.

#### rostrocaudal length

The LC length was measured as the number of axial slices its mask was present in and converted to millimeters.

#### volume

The volume of the LC was determined as the number of voxels in the mask and converted into cubic millimeters.

### 2.4 Experiments

We carried out the following experiments to assess the performance of ELSI-Net in different ways. We compared its segmentations to manual expert ratings and published LC atlases, replicate subject group differences described in the literature, explore correlations of the automatically obtained LC MRI features to CSF biomarkers of AD pathology and assess the influence of acquisition related factors. For the statistical analyses we use Jasp[26] as well as the scipy[27] library.

#### 2.4.1 Mask Similarity

ELSI-Net’s performance is evaluated by determining the agreement of its results with manual subject-wise expert segmentations. We measure the similarity of the masks with the commonly used DSC given by 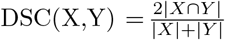 with X and Y being sets of voxels belonging to the respective masks to be compared. Furthermore, we compare the DSC agreement of the fully automated method, ELSI-Net, with a previously published semi-automatic approach that involves manually segmenting the LC on a study-wise template image and transforming the resulting mask into the individual subject spaces. We abbreviate this method with MT and refer the reader to [9] for further details.

#### 2.4.2 Anatomical Agreement

A template-based comparison of ELSI-Net’s LC masks to previously published atlases and the masks of our expert raters allows to assess ELSI-Net’s results with respect to their anatomical plausibility in terms of anatomical position and extent. To this end, we coregistered and morphed the upsampled FLASH scans of all healthy and MCI subjects together with their manual (R1) and ELSI-Net LC segmentations to FSL standard 0.5mm asymmetric MNI space[28] using Advanced Normalization Tools (ANTs)[29] Syn registration with bspline interpolation. We then calculated a probabilistic mask for the ELSI-Net and manual segmentations and binarized both using a 50% threshold. In total, 8 subjects were excluded due to a failure of the registration process. We rendered the ELSI-Net LC template alongside the template obtained from the manual ratings as well as a recently published LC atlas by Dahl and colleagues (so-called meta mask[20]) that combines the information of several established LC atlases[30, 18, 16, 31, 9, 8] and was brought into the same asymmetric MNI space of the other templates using Syn registration. We performed these steps analogously to Dahl et al.[20] The visualization was carried out using 3D Slicer.[32] Additionally, we determine the DSC agreement between the resulting template masks and calculate the agreement with the meta mask by Dahl et al. using the accuracy metric (mean of specificity and sensitivity) as previously described by the authors.[20] The resulting agreement is compared to a number of established and publicly available LC atlases.

#### 2.4.3 Feature-based Comparison of Subject Groups

We conducted significance tests, such as t-Tests (with preceeding Levene tests for equality of variances) and one-way analyses of variance (ANOVAs) as well as Cohen’s d as an effect size measure.

#### 2.4.4 Relationship between LC volume and CSF measures of AD pathology

Measurements of amyloid beta (Aβ_42_/Aβ_40_) and tau proteins (total tau (TTau), phosphorylated tau 181 (PTau)) in the CSF are established biomarkers of Alzheimer’s Disease pathology. From 85 of the 188 DELCODE subjects CSF measures were obtained (AD dementia: 7, mild cognitive impairment: 21, subjective cognitive decline: 22, healthy controls: 35). We correlated the automatically derived LC features to CSF biomarkers of AD pathology using Pearson’s r correlations performed in Jasp.[26]

#### 2.4.5 The Influence of Image Quality and Acquisition Site

We investigate the potential impact of motion artefacts on ELSI-Net. To objectively assess the MRI image quality, we make use of the convolutional neural network based approach to motion artefact quantification as recently proposed.[33] This network was trained to estimate the structural similarity (SSIM) of a single corrupted image slice to its (non-existent) ideal, uncorrupted version. The resulting predicted/estimated SSIM thus quantifies the amount of corruption by motion artefacts, 1.0 encoding perfect image quality and 0.0 the worst. We applied the network to all acquisitions from both datasets by slice-wise processing. We chose the minimum predicted SSIM out of all slices of an acquisition as its image quality score.

Apart from reporting the overall image quality of the used datasets, we investigate the relation of image quality to the DSC agreement between ELSI-Net and the expert’s masks as well as to the disagreement in terms of resulting features (measured as the absolute difference between extracted features such as median and maximum CRs and LC volume of the two segmentation approaches). Finally, we assess the influence of the acquisition site on the agreement between ELSI-Net and the expert rating as well as on the LC CR features per se.

## 3 Results

### 3.1 Mask Similarity

#### 3.1.1 Healthy Aging Dataset

The HAD was rated by both expert raters with an inter-rater agreement of 67.58% ± 8.90% (mean ± standard deviation) DSC when averaging the left and right LC. The left plot in Figure 2 shows that both, our automatic method (73.19% ± 7.75%) as well as the semi-automatic template registration based approach (MT)[9] (74.00% ± 15.56%) perform comparably to the inter-rater agreement in terms of mask similarity when using R1’s segmentations as the reference. However, the values derived using ELSI-Net are subject to a substantially lower standard deviation (almost half of the semi-automated method). ELSI-Net does not show a difference in its agreement to either expert rater, while with MT a strong decline of mask agreement with respect to R2’s segmentations compared to those of R1 is apparent.

**Figure 2:**
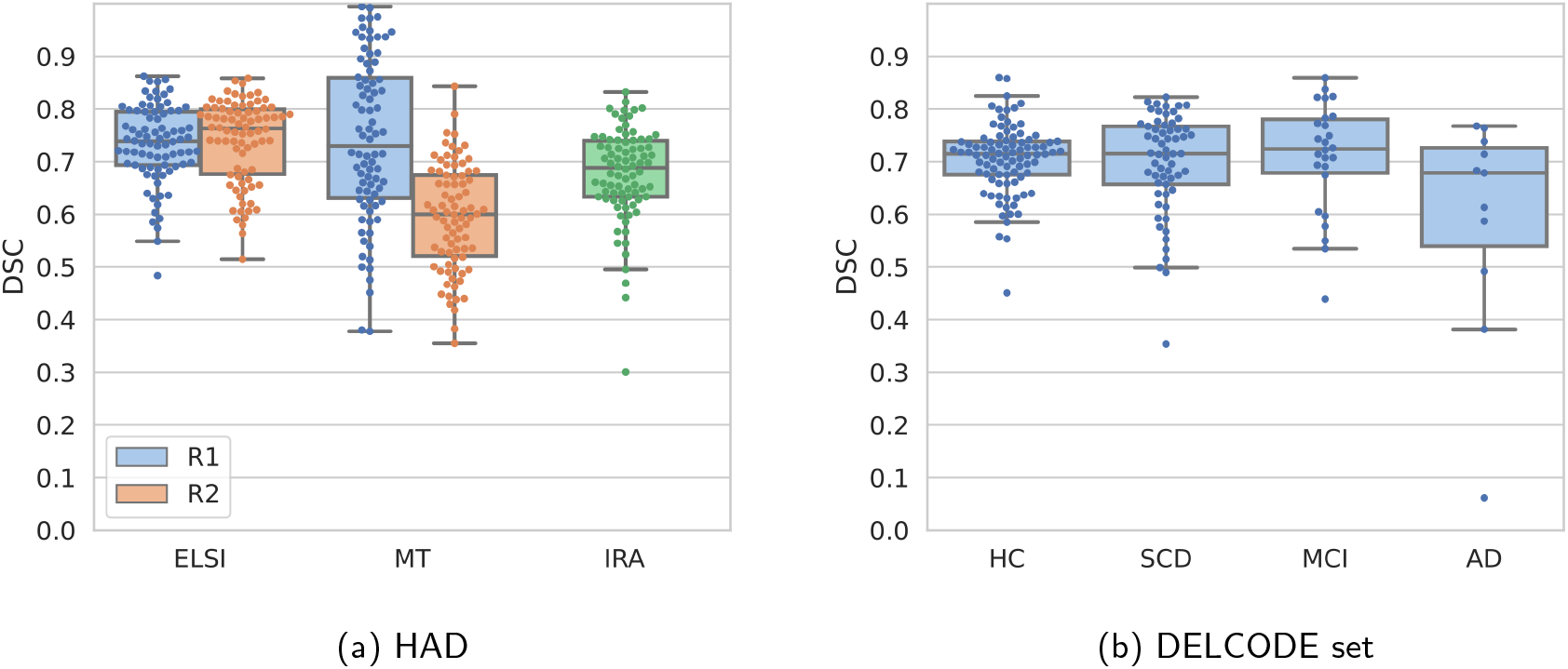
Box- and swarm plots of DSC agreement values of ELSI-Net and the semi-automatic template registration-based method (MT)[9] on Healthy Aging Dataset (a) as well as ELSI-Nets DSC agreement on the DELCODE set (b) with respect to the manual expert segmentations by our experts R1 and R2. They each rated all of HAD, for which we show the agreements to each rater individually (blue and orange hue) and the inter-rater agreement (IRA, green hue). Each expert rated a complimentary subset of the DELCODE study, so that we report the agreement to respective available raters mask.

#### 3.1.2 DELCODE Dataset

Across almost all subject groups from DELCODE, ELSI-Net shows relatively high agreement with a manual expert rating (see Figure 2 b). The mean DSC consistently exceeds 70% and the standard deviations are in a range of 6.4% to 10.2%, which is comparable to the inter-rater agreement measured on HAD. The comparatively small group of AD dementia subjects however constitutes the exception with lower agreement and a larger standard deviation in the DSC values, although the median with 67.86% is in the range of the inter-rater agreement. Figure 3 provides a qualitative visualization of this subject showing the median DSC agreement of the AD dementia group.

**Figure 3:**
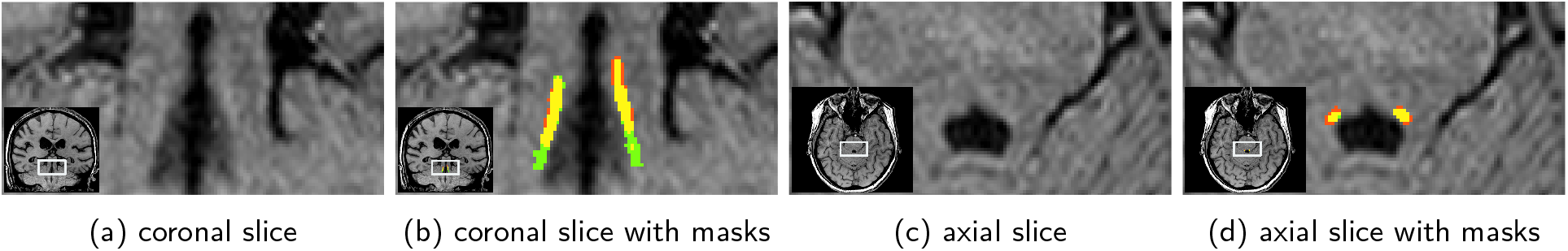
Visualization of an example AD dementia subject from DELCODE with the median DSC agreement from this group (67.86%). In (b) and (d) the segmentations of ELSI-Net (red) are compared to the expert rating (green). Overlap between ELSI-Net and the expert rating is indicated in yellow.

### 3.2 Anatomical Agreement

Figure 4 depicts a template generated from the ELSI-Net results from DELCODE overlaid with a template from the manual ratings as well as a previously published LC atlas (meta mask) from Dahl et al.[20] ELSI-Net’s agreement to the manual rating’s template is very high (92.14% DSC, see Figure 4a). It confirms a good overall agreement to the manual ratings of both raters that was already suggested by the subject-level mask agreement evaluation. Only a very slight discrepancy is visible: the ELSI-Net template appears slightly shifted towards the rostral direction compared to the manual rating’s template. There are however more deviations observable when comparing it to the meta mask (see Figure 4b), which comprises information of multiple other published atlases derived from different MRI acquisitions and modalities. The meta mask is located more rostral than the ELSI-Net template and therefore the template of the manual ratings as well. ELSI-Net’s agreement wrt. the meta mask (64.33% DSC) exceeds the agreement of the manual rating’s template (60.09% DSC) with the meta mask by more than 4% DSC.

**Figure 4:**
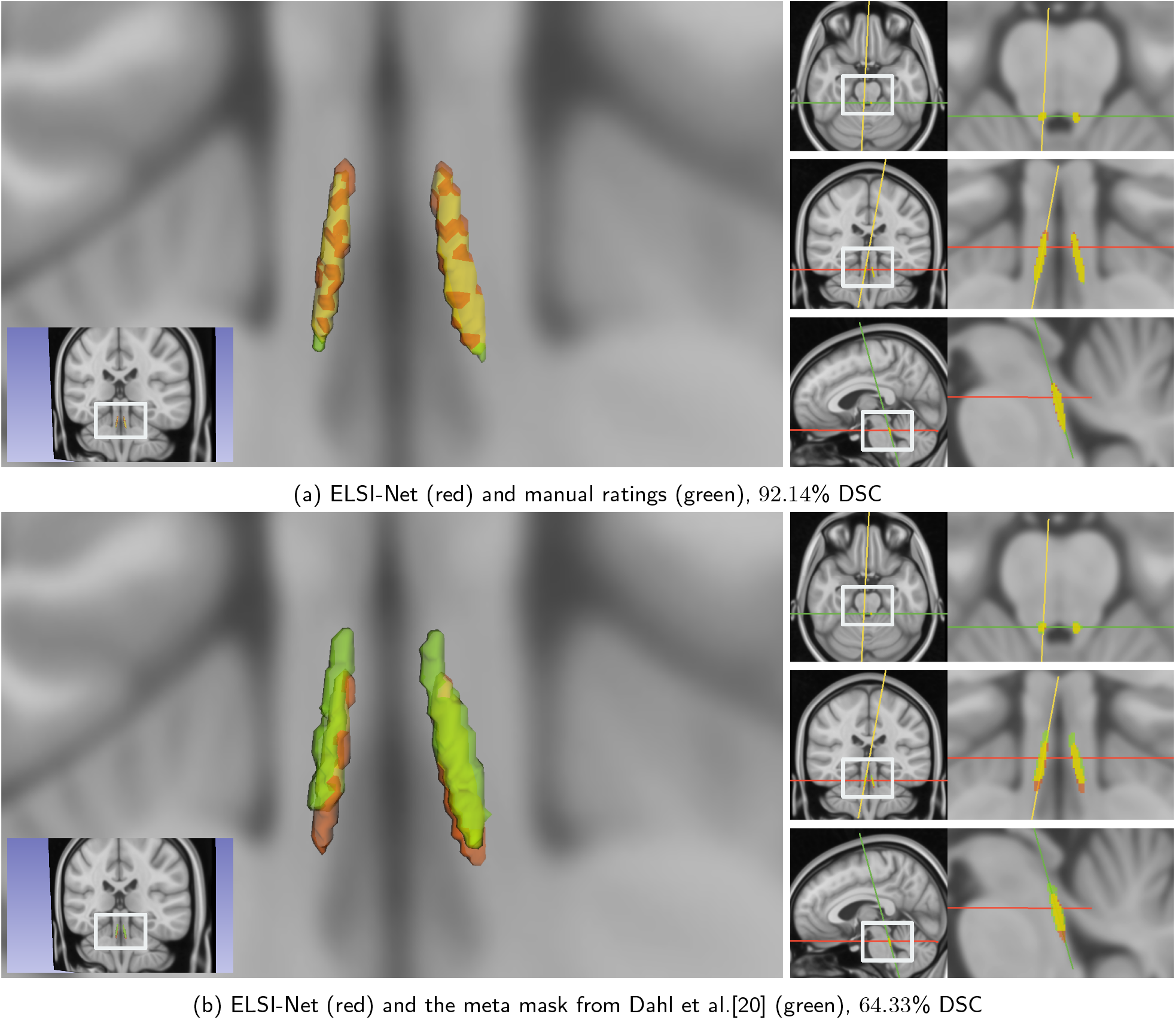
Overlay of the ELSI-Net template from DELCODE with a template generated from manual ratings (see Figure 4a) and a published LC atlas, meta mask, that combines information from several other previously published LC atlases (see Figure 4b) in MNI space (0.5mm). On the left, a 3D rendering of the masks from an example coronal slice overlaid on the MNI template is shown. On the right, the respective axial (top), coronal (middle) and sagittal (bottom) 2D slices are visualized. Red color indicates the ELSI-Net template mask, green color the respective other template and the overlapping volumes are colored in yellow. The corresponding slices are indicated by red (axial), green (coronal) and yellow (sagittal) lines on the right sides, respectively. Mask agreements are provided (DSC). Note that the agreement between the manual template (green in Figure 4a) and the meta mask (green in Figure 4b) is 60.09% DSC.

Other established LC atlases show agreements ranging from 63% to 47% to the meta mask as measured by the accuracy metric specified by the authors[20] (mean of sensitivity and specificity). We calculated the same metric and found ELSI-Net’s template to achieve 66.62% using this measure, which demonstrates a comparatively high agreement with the meta mask. It also exceeds the manual rating’s template score which amounts to 63.86%.

### 3.3 Feature-based Comparison of Subject Groups

We obtained the previously described LC features (see 2.3) from HAD and DELCODE using ELSI-Net and report the resulting distributions here.

#### 3.3.1 Healthy Aging Dataset

We identified significant differences in LC features between young and older subject groups. The maximum CR and particularly the rostral maximum subregional CR halves and thirds show age related increases in LC intensity on HAD. Figure 5 visualizes the resulting distributions for these features and groups. We determined Cohen’s d for estimating the effect size of the difference between young and older subjects and found that with maximum CRs R1, R2 and ELSI-Net resulted in 0.489, 0.596 and 0.662, respectively. Furthermore, all of these differences in the maximum CR were found to be statistically significant (e.g. ELSI-Net maximum CR: Student t-Test(2.759, p = 0.007), Levene test for equality of variances (1.857, p = 0.177)). When inspecting the subregional LC CRs, it becomes evident that the age related effect appears to be stronger in the medial and rostral LC parts. We find an increasing effect size measured by Cohen’s d from 0.374 (caudal) to 0.489 (medial) and 0.705 (rostral) for the maximum subregional CRs (splitting LC in thirds of equal length) determined by ELSI-Net.

**Figure 5:**
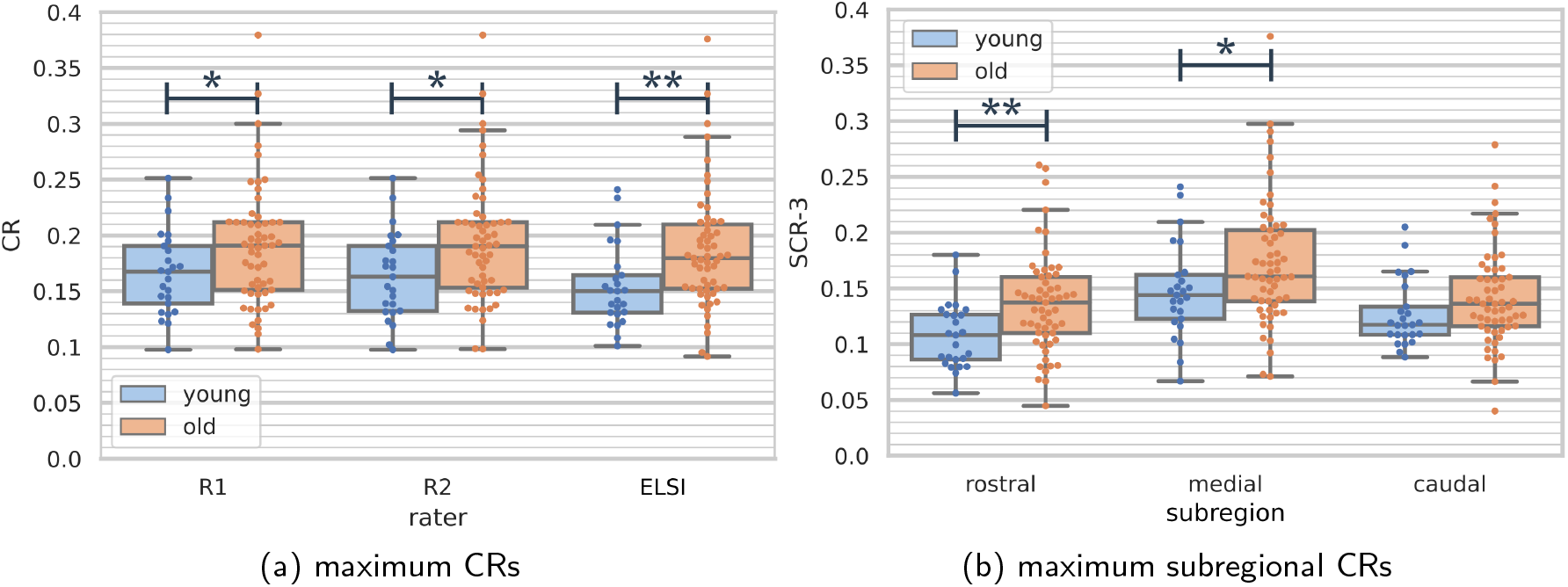
Box- and swarm plots of maximum LC CRs (a) and subregional maximum LC CRs (b) in young (blue) and older (orange) subject groups from Healthy Aging Dataset. For a) the values of the expert raters R1 and R2 are reported as well as those of the fully automatic ELSI-Net. In plot (b) only the ELSI-Net results are shown. Significant differences are indicated by *(p < 0.05) and **(p < 0.01) using a two-tailed t-Test.

#### 3.3.2 DELCODE set

Figure 6 visualizes the subject group distributions of four LC features obtained with ELSI-Net and compares them to the manual rating’s distributions. A trend of decreasing LC volume and length as determined by ELSI-Net appears with increasing clinical severity of the subject groups, which is not present in the expert’s LC volume and length measurements. The visible decrease in volume in the AD dementia group (see Figure 6c) bears resemblance to the trend measured by the median CR with the expert ratings. A one-way ANOVA confirmed the statistical significance of this decrease (healthy controls vs. AD dementia: F = 5.258, p = 0.002 and post-hoc p_Tukey_ = 0.006). We found Cohen’s d for the difference in volume between the healthy group and the AD dementia subjects to be 1.089 with ELSI-Net, which is comparable to the effect size found with the manual rating and the median CR group difference (Cohen’s d: 0.993, one-way ANOVA: F = 3.575, p = 0.015 and post-hoc p_Tukey_ = 0.016). Similar to LC volume, the LC length feature measured by ELSI-Net was also significantly decreased in AD dementia(one-way ANOVA: 4.084, p = 0.008 and post-hoc p_Tukey_ = 0.009, Cohen’s d: 1.036))(see Figure 6d). The group differences in the CRs using ELSI-Net were not impacted by the choice of the reference region generation approach. Significant group differences were neither identified with the semi-automatically nor with the automatically generated reference regions.

**Figure 6:**
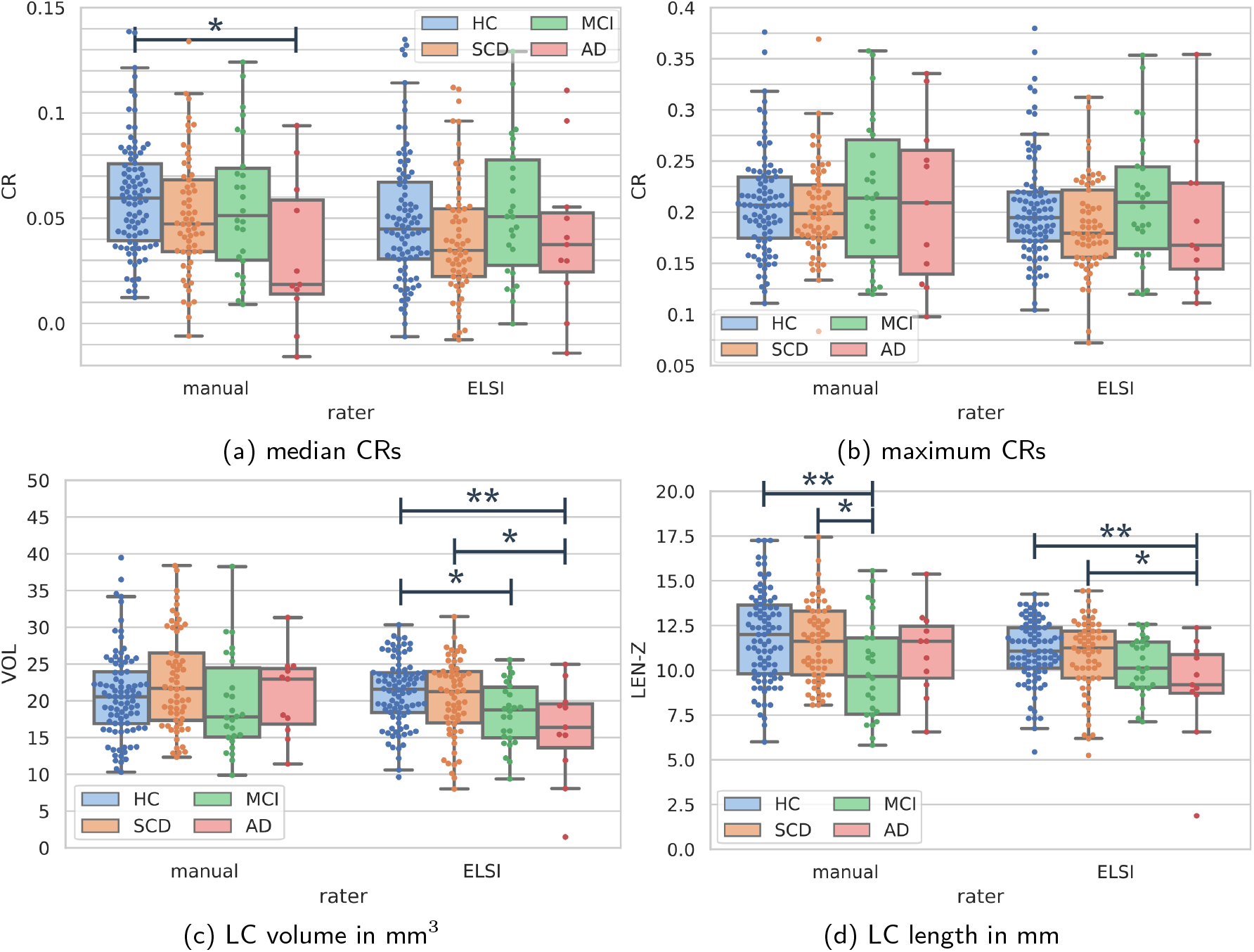
Box- and swarm plots of selected features (median LC CRs (a), maximum CR (b), LC volume (c), LC length (d)) of the manual rating (left) and ELSI-Net (right) on the healthy (blue), subjective cognitive decline (orange), MCI (green) and AD dementia (red) subject groups of the DELCODE set. Significant differences are indicated by *(p < 0.05) and **(p < 0.01) encoding the Tukey post-hoc test result of the respective one-way ANOVA (with p < 0.05).

### 3.4 Relationship between LC volume and CSF measures of AD pathology

Motivated by the observed decreases in LC volume in MCI and AD dementia subjects measured by ELSI-Net, we correlate this feature to all available CSF measures of AD pathology. Several significant correlation results were found. They are reported in Figure 7. Although the correlations are weak, they show decreased LC volume is associated with higher tau and amyloid pathology. We found similar correlations between LC length and amyloid pathology (LC length and Aβ_42_/Aβ_40_: r = 0.228, p = 0.036; LC length and Aβ_42_/PTau181: r = 0.298, p = 0.006), but no correlation with LC CR features.

**Figure 7:**
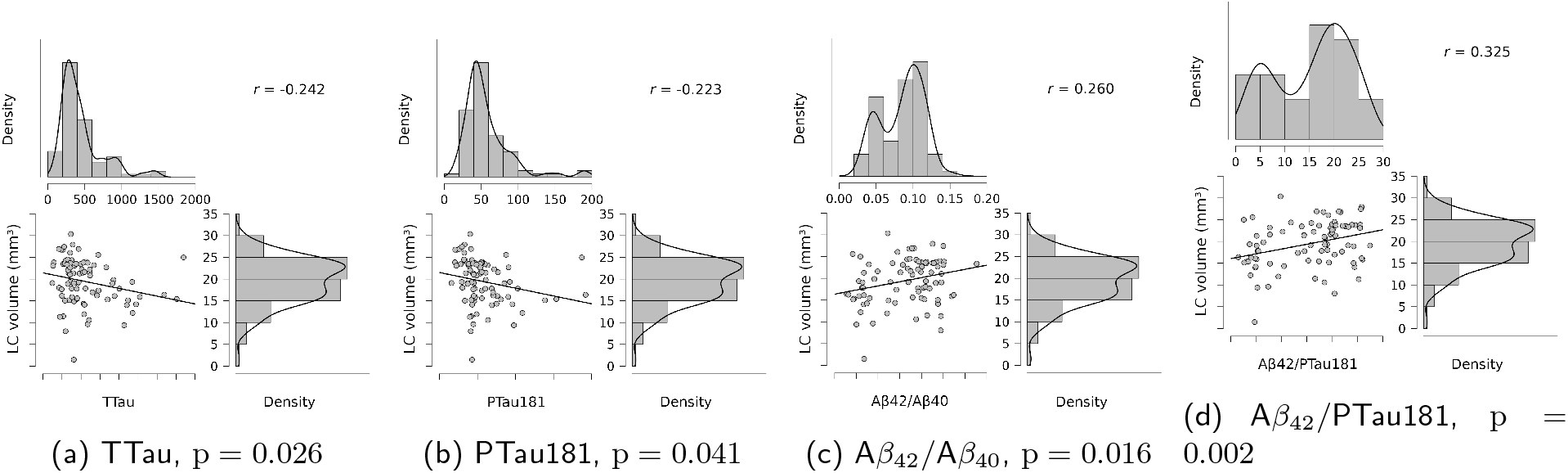
Pearson’s r correlations between LC volume obtained with ELSI-Net and CSF measures of AD pathology. Scatterplots and value distributions are visualized. Correlation coefficient (r) and p-value are reported.

### 3.5 The Influence of Image Quality and Acquisition Site

Using a recently proposed method for the quantification of motion artefacts,[33] we conducted the previously described experiments to estimate the influence of acquisition artefacts on the LC metrics.

#### 3.5.1 Image Quality of the Datasets

When comparing the image quality of the two datasets, it becomes apparent that the DELCODE dataset (SSIM mean: 0.794, standard deviation: 0.058) shows lower quality than HAD (SSIM mean: 0.827, standard deviation: 0.035) (Welch t-Test: −5.874, p = 1.394E − 8, Levene test for equality of variances (11.505, p = 7.987E − 4)). A one-way ANOVA showed no significant differences between DELCODE subject groups (1.629, p = 0.184). Nonetheless, a coincidence of slightly decreasing image quality with increasing clinical severity is immanent in our particular dataset. The SSIM means of the healthy control, subjective cognitive decline, MCI and AD dementia groups are 0.795, 0.802, 0.784 and 0.764, respectively. This motivates further investigation of a potential influence of image quality on subject group differences.

#### 3.5.2 Correlation of Image Quality with Segmentation Performance and Feature Deviation

We computed several image quality related Pearson’s r correlations that are reported in Table 1. One of them is between ELSI-Net’s mask similarity to the manual expert rating on DELCODE quantified in terms of DSC and the measured image quality (predicted SSIM) of the samples. Although the correlation is significant it does not appear strong (r = 0.239, p < 0.001). However, a modest correlation between image quality and absolute differences in maximum CR between the two segmentation approaches was observed (r = −0.401, p < 0.001). It indicates a correlation between image quality and agreement between the CRs of ELSI-Net and manual ratings so that with increasing image quality, there is greater agreement between the two segmentation approaches. This correlation is much weaker for median LC CRs (r = −0.255, p < 0.001) and was not observed for LC volume feature extraction indicating LC volume may be influenced less by image quality.

**Table 1:**
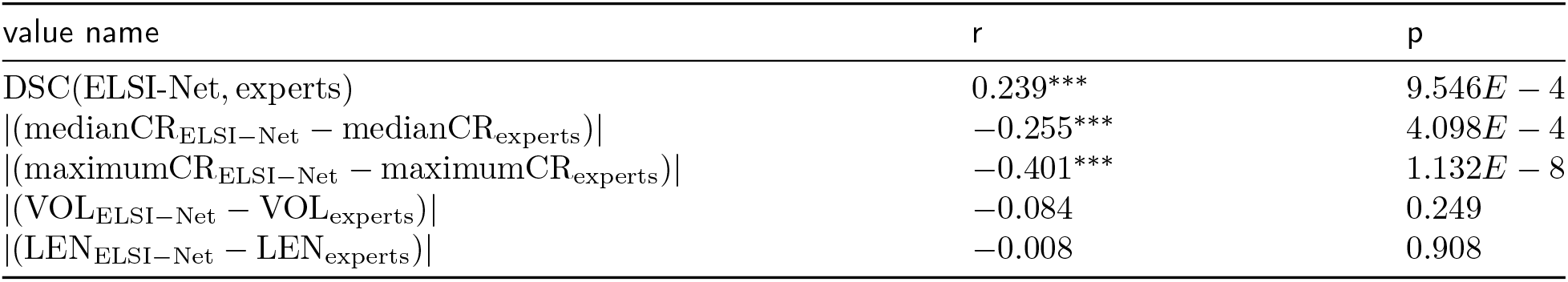
Pearson’s r correlation of the image quality measured as predicted SSIM of DELCODE and the specified other values: the DSC agreement of the ELSI-Net and expert’s masks and the absolute differences of several extracted LC features. ^*^ p < 0.05, ^**^ p < 0.01, ^***^ p < 0.001

#### 3.5.3 Assessment of Acquisition Site Effects

Several one-way ANOVAs were carried out on DELCODE to gain insights into the potential influence of the acquisition sites. We found significant image quality (measured by predicted SSIM) differences between sites (one-way ANOVA: 13.274, p = 4.121 × 10^−6^, one of the four sites was excluded since it contributed only one of the subjects). By means of further one-way ANOVAs, we found no significant site effects on mask agreement between ELSI-Net and the expert rating (measured by DSC) as well as agreement on median and maximum CRs. This indicates that the agreement between ELSI-Net and manual expert ratings is not affected by the acquisition site. However, site related differences could be identified in all of the CR and subregional CR features themselves (e.g. median CR, one-way ANOVA 26.809, p = 6.057 × 10^−11^). In contrast, ELSI-Net estimates of LC volume and length are not subject to site effects.

## 4 Discussion

Here, we report an improved fully automatic segmentation and feature extraction method for in vivo assessment of the LC. This method comprises an ensemble approach to applying the neural networks for LC localization and segmentation. Following the initial localization, an additional intensity normalization step using a local patch surrounding the LC is applied. We propose a method for the generation of a reference region removing the need for an additional segmentation of the Pons region. This approach is validated in healthy young and older adults as well as in adults on the AD dementia spectrum for robustness with respect to previously published LC segmentation approaches and clinical observations. The proposed changes, most notably the addition of the ensemble-based inferences led to performance improvements in terms of DSC agreement compared to our previous work[14], where the same nested cross-validation evaluation scheme, data splits, and dataset (HAD) were used.

In healthy aging, ELSI-Net was able to segment the LC with very high accuracy performing equal or better than an expert manual rater. In contrast to a previously published semi-automatic approach,[9] ELSI-Net exceeds the inter-rater agreement in terms of DSC with respect to both raters (compare ‘ELSI’ and ‘MT’ in Fig. 2a). It is therefore arguably more objective than both the semi-automatic segmentation method and a single expert rater. A higher Cohen’s d value with respect to age-related differences between young and older adults shows that ELSI-Net could detect these differences reliably and with increased sensitivity compared to a manual segmentation approach. ELSI-Net was also able to replicate previously observed age-related increases in rostral and middle LC contrast using the same dataset.[9] ELSI-Net may potentially be deployed across sites to further reduce rater bias of LC analyses and increase comparability of studies using similar subjects and MRI protocols, e. g. to determine normative feature ranges of a healthy LC given a specific age. Indeed, acquisition site related influences on the LC features were found independently of the segmentation approach and have to be considered.

In a clinical cohort of individuals with AD dementia from DELCODE, ELSI-Net could effectively segment the LC without any fine-tuning, solely being trained on the healthy aging dataset. For the most part (including the MCI subject group) a satisfactory performance measured by DSC compared to the experts’ rating was achieved. The AD dementia group was a noticeable exception, although it was the smallest group with only 11 participants and further analyses are required in larger cohorts of individuals with AD dementia to comprehensively determine its segmentation accuracy.

The overall anatomical plausibility of the automatically obtained LC masks and a performance comparable to those of experts is indicated by the very high agreement between the LC template of the ELSI-Net DELCODE masks and the template generated from the manual ratings, but also a meta mask comprising a number of previously published atlases.[20] ELSI-Net’s template shows the highest agreement with the meta mask among all published atlases that were used for the meta mask creation. This indicates that ELSI-Net can generate anatomically precise LC segmentations removing the need for semi-automatic or manual segmentation. With ELSI-Net we found significant differences between healthy controls and subjects with AD dementia with respect to LC volume and length but not median LC contrast, as observed with the expert ratings. This could indicate a deviation of segmentation style between ELSI-Net and manual LC segmentation. It is conceivable that ELSI-Net is differentially influenced by a reduction in LC MRI contrast present in AD dementia subjects compared to expert raters that rely more on anatomical prior knowledge. ELSI-Net was only trained on LC segmentations from healthy young and older adults. Since it has never seen the variance introduced by AD dementia during training, the lower LC contrast or additional data characteristics unknown to us may yield smaller LCs in ELSI-Net’s assessment in AD. This would explain the lower DSC agreement between ELSI-Net and manual raters observed in the AD dementia group as well as the deviations seen in LC CRs between ELSI-Net and manual rating on DELCODE. Therefore LC volume and length estimates using ELSI-Net might be more accurate in clinical cohorts due to reduced human bias that manual raters may be prone to. It should be noted that ELSI-Net may not only rely on intensity, but on additional characteristics such as the surrounding anatomy and shape of the LC during the segmentation procedure.

We further investigated the influence of image quality on our (automatic) LC analysis in multiple ways. We found significant correlations between image quality and agreement on CR-based features (as quantified by absolute difference between ELSI-Net and the experts rating in particular for maximum intensity CRs). However, no such correlation was observed with respect to ELSI-Net’s LC volume measure, where we observed most pronounced differences between healthy controls and AD dementia. We found significant acquisition site related effects on the image quality and CR features in general. No influence of site could be found on the agreement between ELSI-Net and the experts’ rating with respect to mask and CR feature agreement as well as ELSI-Net’s measurements of LC volume and length indicating a robust performance across multiple sites. As an additional validation step, we further assessed how LC segmentations generated by ELSI-Net are related to previously reported associations with AD pathology. In a subset of subjects with known CSF status from DELCODE, we found reduced LC volume was significantly associated with increased tau and amyloid pathology in agreement with previous findings.[10, 34, 35, 36]

### 4.1 Limitations

An important caveat in our analyses is the rather small number of AD dementia subjects (n = 11) in our clinical cohort of 188 participants. Further evaluations on more datasets with larger groups of AD dementia subjects, ideally with amyloid and tau pathology biomarkers, are required to ascertain the performance of ELSI-Net more conclusively in this population. Another interesting aspect that was left unexplored is the robustness of ELSI-Net with respect to acquisition parameters and differing MRI protocols. Both of the cohorts investigated here comprised T_1_-weighted FLASH imaging, hence it remains to be explored how ELSI-Net performs on more diverse datasets using alternative LC MR imaging modalities. Of course, a necessary requirement for the application of ELSI-Net is the acquisition of specific sequences with LC contrast in general. Recently, automatic methods were applied to LC segmentation in acquisitions without LC contrast as an alternative to atlas-based approaches.[37] While inherently lacking precision and the possibility to extract LC contrast, volume or length features, they may allow functional MRI or diffusion MRI analyses in datasets where LC MRI was not acquired.

Finally, the image quality assessed here was based on the quality of the whole-brain acquisition and was not LC specific, which may not reflect the quality of the visualization of the LC and its immediate vicinity.

### 4.2 Conclusion

In this work, we evaluate an improved version of a previously proposed, fully automatic approach to LC segmentation and feature extraction termed ELSI-Net. Evaluation on LC imaging data acquired from young and older adults but also from subjects across the AD dementia continuum show that ELSI-Net reliably generates anatomically plausible results with excellent agreement to established LC atlases. We found increased objectivity with ELSI-Net compared to single expert raters and a semi-automatic LC segmentation method. LC features by ELSI-Net demonstrate high sensitivity replicating previously shown subject group differences in healthy aging and AD dementia. We saw correlations of LC volume measured by ELSI-Net to tau and amyloid pathology and robust performance with respect to data acquired across multiple sites.

ELSI-Net provides a means to automatically segment the LC with high accuracy particularly in aging cohorts. Further analyses are required to determine its effectiveness in segmenting the LC in different LC MRI contrasts, longitudinal datasets and additional clinical cohorts e.g. including subjects with Parkinson’s Disease, depression and further neurological disorders.

## Acknowledgments

None.

